# Research Synergy and Drug Development: Bright Stars in Neighboring Constellations

**DOI:** 10.1101/149559

**Authors:** Samet Keserci, Eric B. Livingston, Lingtian Wan, Alexander R. Pico, George Chacko

**Affiliations:** NETE Labs, NET ESolutions Corporation, McLean, VA, USA; Research Intelligence, Elsevier Inc., New York, NY, USA; Gladstone Institutes, San Francisco, CA, USA

## Abstract

Drug discovery and subsequent availability of a new breakthrough therapeutic or ‘cure’ is a compelling example of societal benefit from research advances. These advances are invariably collaborative, involving the contributions of many scientists to a discovery network in which theory and experiment are built upon. To understand such scientific advances, data mining of public and commercial data sources coupled with network analysis can be used as a digital methodology to assemble and analyze component events in the history of a therapeutic. This methodology is extensible beyond the history of therapeutics and its use more generally supports (i) efficiency in exploring the scientific history of a research advance (ii) documenting and understanding collaboration (iii) portfolio analysis, planning and optimization (iv) communication of the societal value of research. As a proof of principle, we have conducted a case study of five anti-cancer therapeutics. We have linked the work of roughly 237,000 authors in 106,000 scientific publications that capture the research crucial for the development of these five therapeutics. We have enriched the content of networks of these therapeutics by annotating them with information on research awards as well as peer review that preceded these awards. Applying retrospective citation discovery, we have identified a core set of publications cited in the networks of all five therapeutics and additional intersections in combinations of networks as well as awards from the National Institutes of Health that supported this research. Lastly, we have mapped these awards to their cognate peer review panels, identifying another layer of collaborative scientific activity that influenced the research represented in these networks.

## Introduction

Data mining of public data sources coupled with network analysis enables the quantitative description of research discoveries that were influential in the development of a breakthrough therapeutic or ‘cure’. The set of scientific publications, clinical trials, patents, and regulatory approvals, linked to each other by citation or assignment, that documents the progress of concepts from basic research to a cure is termed a ‘cure network’ [1]. Reconstructing such networks enables a deeper contextual understanding of knowledge diffusion across disciplines, scientific interests, culture, and time [2]. Such studies also (i) provide evidence for the broad collaborative platform of basic and translational research underlying major scientific advances such as cures for diseases [3] (ii) support strategic communications to oversight bodies, and (iii) help communicate the societal value of research. The understanding of a therapeutic network, when coupled with information from clinical use of a therapeutic, also enables a recursive learning of the pathogenesis of the disease it is being used to treat, as has been noted for the burgeoning field of immunotherapeutics [4].

Williams and colleagues have elegantly demonstrated the feasibility and value of data mining and network analysis using, as case studies, ivacaftor and ipilimumab, approved for the treatment of cystic fibrosis and melanoma respectively (*vide supra*). They observed that ‘the nature of a cure discovery network is complex and fundamentally collaborative’, noting in the case of ivacaftor, that at least 7,067 scientists with 5,666 unique affiliations contributed to ivacaftor-relevant research over a period greater than 100 years. These authors also suggest that thoughtful metrics derived from this concept could inform decision making by funders.

Extending the methodology to study additional cures and significant research advances is a logical next step. Ascertaining the nature of the interactions, if any, between networks, is also of considerable interest since it supports an understanding of collaboration across networks as well as common features of science networks. Lastly, even if ambitious, scaling from case studies to mapping the entire domain of drug development is likely to be beneficial in planning, resource allocation and optimization of drug development activities.

To address these questions, we have built upon prior art for single networks to (i) incorporate enhanced data mining methods and network metrics (ii) include enriched data from a commercially available bibliographic database with disambiguated author identifiers (iii) include information on research awards and peer review of grants (iv) extended single network analysis to map publications and authors across multiple networks. Key assumptions in constructing these networks are that the references found in relevant documents are appropriate citations of new knowledge relevant to a given cure and that a further retrospective round of citation discovery will reveal previous influential discovery [1]. For evaluation, we have conducted case studies of a cluster of five FDA-approved therapeutics for cancer. We present the results of this case study as a body of work for further study by other researchers, and a step towards mapping the universe of FDA-approved drugs and biologicals.

## Materials and Methods

A set of five anti-cancer therapeutics, three drugs and two biologicals, approved for use in humans by the Food and Drug Administration (FDA) was selected for this study (Table 1). Imatinib [5] and Sunitinib [6] are tyrosine kinase inhibitors, Nelarabine [7] is a nucleoside analog, and Ramucirumab [8] and Alemtuzumab (Campath) [9] are humanized antibodies that target the CD52 and vEGFR-2 cell surface receptors respectively. For each of these therapeutics, a set of relevant scientific publications was constructed as in Williams et al. [1] but with specific modifications detailed below. An allowance of 2 months was also made for ‘publication lag’ when assembling referenced material. For example, if a therapeutic was approved on Jan 1, 2017, documents published on or before March 31, 2017 were included. For each of the five therapeutics, a first-generation list of PubMed identifiers (citing_pmid) was harvested from the five different data sources (Table 1).

**Table 1.**
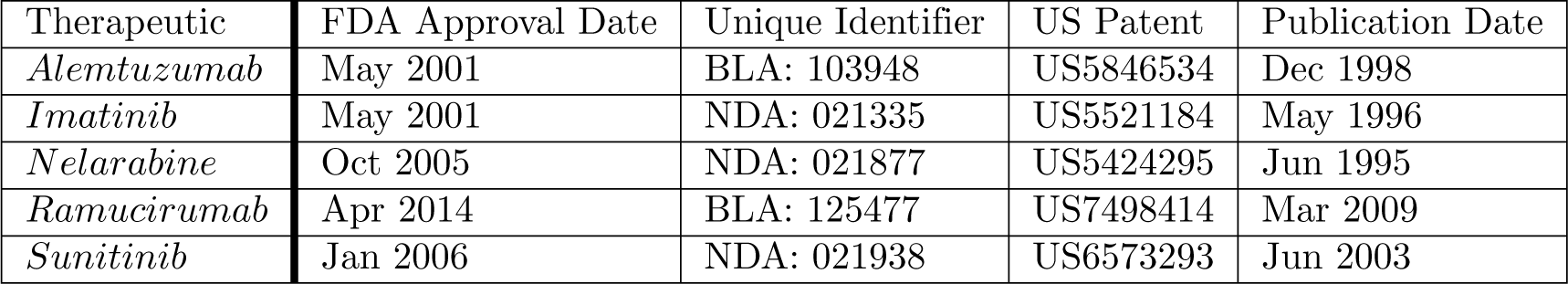
Case Studies of Five Anti-Cancer Agents. Five anti-cancer therapeutics, with FDA approval dates ranging from 2001 to 2014, were selected as case studies. The unique identifier for each therapeutic is an FDA assigned NDA or BLA number. While multiple patents are typically associated with a drug or biological, the single US patent number displayed represents the primary invention that preceded approval of the therapeutic. The publication date for each patent is listed in the last column.

| Therapeutic | FDA Approval Date | Unique Identifier | US Patent | Publication Date |
| --- | --- | --- | --- | --- |
| *Alemtuzumab* | May 2001 | BLA: 103948 | US5846534 | Dec 1998 |
| *Imatinib* | May 2001 | NDA: 021335 | US5521184 | May 1996 |
| *Nelarabine* | Oct 2005 | NDA: 021877 | US5424295 | Jun 1995 |
| *Ramucirumab* | Apr 2014 | BLA: 125477 | US7498414 | Mar 2009 |
| *Sunitinib* | Jan 2006 | NDA: 021938 | US6573293 | Jun 2003 |

### Clinical trials

The national clinical trials database (clinicaltrials.gov) was searched for clinical trials of the five therapeutics that completed by the data of FDA approval. Both cited references and publications from these clinical trials were collected if they were published within the approval date plus two months. To capture publications associated with the clinical trials that were not displayed in clinical trials.gov, PubMed was also searched with the unique identifier (NCT number) of any clinical trials that were identified. To capture publications of clinical trials not registered in clinicaltrials.gov, PubMed was searched using the therapeutic name as keyword, publication type as “clinical trial”, and an appropriate date restriction as in searches of clinicaltrials.gov. For example, the search term (((“alemtuzumab” [Supplementary Concept] OR “alemtuzumab” [All Fields]) OR (“alemtuzumab” [Supplementary Concept] OR “alemtuzumab” [All Fields] OR “campath” [All Fields])) AND (“1900/0101” [PDAT]: “2001/07/31” [PDAT])) AND “clinical trial” [Publication Type] was used to identify publications of clinical trials for Alemtuzumab.

### FDA documents

The drugs@fda website [10] was searched for each of the five therapeutics and cited references in the medical review document were manually extracted and matched to pmids. FDA Approval Summaries published in journals by FDA staff, were available for all five therapeutics and contain cited references. If the published date of a cited reference in an Approval Summary exceeded the approval date plus two months, the publication was not included.

### Patents

For each therapeutic a single patent was identified that best represented the most relevant invention to the therapeutic at hand. Identification of this patent was performed using multiple web sources. The US patent number was then used to identify the patent and the non-patent citation list from Google Patents [11] was manually processed by searching PubMed for appropriate pmids. The accuracy of manual searches was far higher than a citation matching tool that we developed for for this purpose,and were used to generate the data in this study.

### Post-approval literature reviews

Review articles published after a therapeutic’s approval by the FDA are independent studies of the the development of a therapeutic. Accordingly, PubMed was searched for review articles on these five therapeutics that were published between the date of FDA approval and a year following the date of approval. Cited references in these reviews were extracted using PubMed and Scopus. The review articles themselves were not included.

### Pre-approval literature searches

Literature searches were performed using PubMed with a date range of 1900/01/01 to two months post-FDA approval. For example, the search term ((alemtuzumab) OR campath) AND (“1900/01/01” [Date - Publication]: “2001/07/31” [Date - Publication]) was used to retrieve articles of interest relevant to alemtuzumab.

### PubMed and Scopus

Citing_pmids from the five different sources above were combined and deduplicated. Using the Scopus database and its APIs. The manually-generated list of pmids taken from the five sources mentioned (Clinical Trials, FDA Documents, Patents, Post-Approval Literature, and Pre-Approval Literature Searches) were searched in Scopus, using the basic Scopus Search API, to arrive at a list of Scopus IDs (citing_sid) for the publications. The Scopus Abstract Retrieval API was then used to retrieve a more comprehensive record for each of the SIDs comprising that list of publications Next, for each of these publication records citing_sid), we used the Scopus Author Retrieval API to retrieve a full record for each unique author in the publication set. We also used the Abstract Retrieval API to collect records for each of the publications cited by the first generation of publications. This set of cited publications is the cited_sid set. Using the same Author Retrieval API, we then gathered data for each of the unique authors affiliated with the cited_sid publications. Completion of the process yields two sets of publications, citing_sid and cited_sid, with citation links between them and full information on all authors for both generations. Finally, for each author in the study, we used the standard Scopus Search API once more to retrieve a smaller record for every publication affiliated with them in Scopus, in order to tally their overall publication output. While author records in Scopus have overall publication counts as part of the record, by manually downloading each of them, we can store and count them by type (i.e. article, book chapter, Editorial, review, etc.). This allowed us to more precisely arrive at publication totals for only those publication types that are relevant for this study.

Whereas mapping between PubMed and Scopus identifiers at the citing_pmid and citing_sid stage resulted in 1% or less information loss, mapping at the cited_sid to cited_pmid resulted in a loss of roughly 15-20 % of target records. Accordingly the Scopus data was used as the backbone of the publication component of the network and the cited_pmid information was treated as an annotation layer. These observations are summarized in Table 2.

**Table 2.** Citation Counts and Mapping Between Bibliographic Databases. Five anti-cancer therapeutics were selected as case studies. A foundational set of references (citing_pmid) was assembled for each therapeutic from patents, clinical trials, regulatory documents, and the scientific literature (Materials and Methods). Citing_pmids were mapped to Scopus identifiers (citing_sid), which were used, in turn, to retrieve cited publications (cited_sid). Cited_sids were mapped back to PubMed identifiers (cited_pmid). The number of identifiers at each stage of the mapping process is shown along with percentage loss (in parentheses) when mapping across PubMed and Scopus or due to null values in the cited_sid field

| Therapeutic | citing_pmid count | citing_sid count | cited_sid count | cited_pmid count |
| --- | --- | --- | --- | --- |
| *Alemtuzumab* | 599 | 587 (1%) | 8840 (2%) | 7071(20%) |
| *Imatinib* | 1380 | 1373(1%) | 27326(1%) | 23340(17%) |
| *Nelarabine* | 104 | 104(0%) | 2476(1%) | 1990(20%) |
| *Ramucirumab* | 1820 | 1804(1%) | 48587(0%) | 40973(19%) |
| *Sunitinib* | 1512 | 1509(0%) | 33895(0%) | 28661(15%) |

Both citing and cited pmids were mapped to NIH grants and peer review panels (study sections) using public information available through NIHExPORTER [12]. Thus, we enriched our network data by identifying those study sections associated with the awards that supported publications in our networks.

### Networks

The resultant data were modeled as networks and analyzed using metrics based on network topology. We calculated the propagated in-degree rank (PIR) and ratio of basic rankings (RBR) metrics of Williams [1]. PIR represents the sum of aggregated citation scores (first and second degree only) for all articles in a network that can be attributed to an author. In addition to computing PIR for all authors in each network, we also combined the citation data for all five networks and computed a networkPIR (nPIR) score, which was also normalized to the sum of individual PIR scores within each network as the PIRpartitionRatio (PPR) as a way to measure inter-network influence. RBR is intended to represent the fraction of a researcher’s output that is in a network and is defined as the ratio of the number of publications in network to the number of publications in a background dataset for an author. In its original specification, the background dataset for RBR was constructed by keyword searches of PubMed. A potential weakness of this keyword based approach is that keywords do not effectively capture the field or the total output of an author even if multiple background samples are taken. Therefore, we created two new variants of the RBR; network RBR (nRBR) and global-based RBR (gRBR). nRBR uses all publications in our set of five therapeutics as background and gRBR takes advantage of the Scopus author_id to capture the total article output of an author as background. Thus, nRBR and gRBR normalize a researcher’s in-network contributions to backgrounds based on total network and total researcher productivity respectively. The details of how these metrics were calculated are provided in supplementary material S1 File.

### Analysis

All data used in this study were acquired exclusively from the sources listed above. Computations were performed on infrastructure owned or leased by NET ESolutions, Elsevier, or the Gladstone Institutes. Code and scripts used in this study were written in Java, Python, and R and are archived on a publicly accessible Github repository [13]. Network visualization was performed using Cytoscape [14].

## Results and Discussion

### Publications

Scientific publications form the backbone of each of these five networks. Our initial assumptions of appropriate citation and retrospective citation discovery (Introduction) suggest that network nodes that are common to multiple networks are likely to be influential. We calculated intersection counts for all possible combinations of publications in the Alemtuzumab, Imatinib, Nelarabine, Ramucirumab, and Sunitinib networks (S2 Table 1). We also applied intersection analysis at a finer level of granularity by computing intersection counts for both first generation citations (citing_pmid) and second generation citations (cited_sid). The results are displayed as Venn diagrams in Fig. 1.

**Fig 1.**
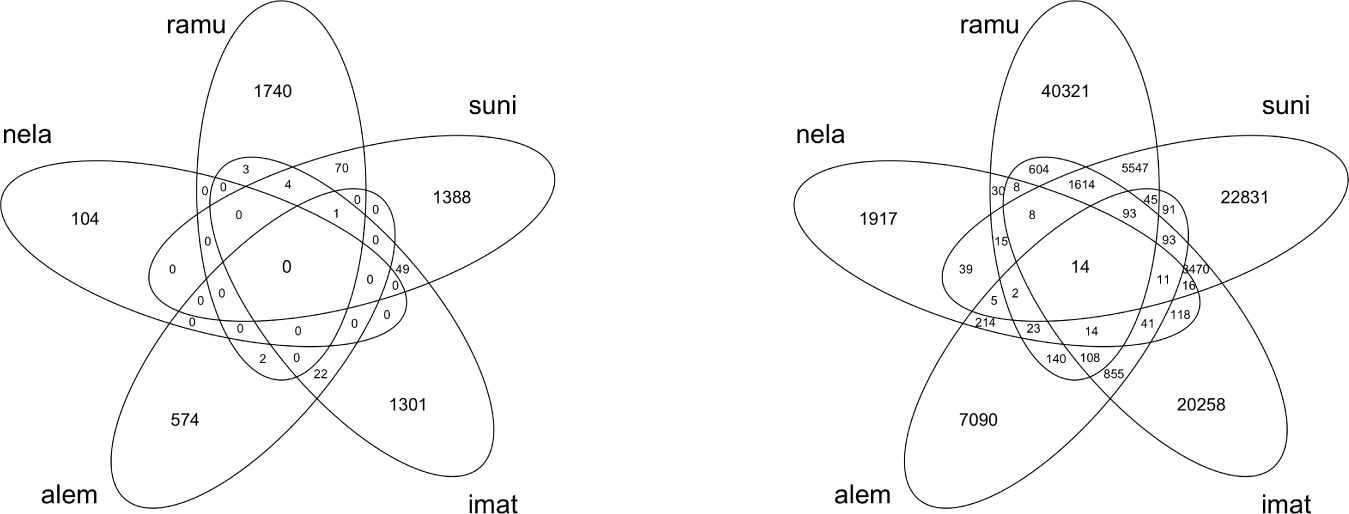
Intersecting Publications in Five Networks. Intersections were calculated across all five networks for the first generation of references (citing_pmids) and as well as for the second generation of references (cited_sids) and displayed as Venn diagrams. *Left Panel.* No first generation publications are observed common to all five networks. A single publication is cited in four of five networks. *Right Panel.* 14 publications are common to all five networks. Abbreviations: alem (Alemtuzumab), imat (Imatinib), nela (Nelarabine), ramu (Ramucirumab), suni(Sunitinib)

The intersection of all five networks consists of 14 publications out of a total of 106,720 unique Scopus identifiers (Supplemental Data, Table 1). Strikingly, not even a single publication is common to all five networks at the first generation level (citing_sid) although a single publication, the pathbreaking work of Kohler and Milstein on the production of monoclonal antibodies [15], is cited in four out of five networks. All 14 publications are in the second generation of citations (cited_sid) and another 198 comprise the sum of intersections in all possible four-network combinations, roughly an order of magnitude greater than the case of cited references. We manually grouped these 14 publications using high level descriptive terms and observed that this group was composed of statistical methods (5 publications), molecular and cell biological methods (4 publications), analytical and structural biology techniques (3 publications) and cancer biology (2 publications). Of these last two, one is a review of the p53 gene [16] and the second is a study of angiogenesis in children with acute lymphoblastic leukemia [17]. Thus, the majority of this small set of 14 publications describes methods that are heavily cited in these therapeutic development networks, which is consistent with observations of the general scientific literature [18]. The relationship between core publications and their therapeutic networks is visualized in Fig. 2. As the subject of another study, we are actively working on a scalable automated strategy to characterize the entire dataset as well as all combinations of intersections between networks using high level descriptive terms.

**Fig 2.**
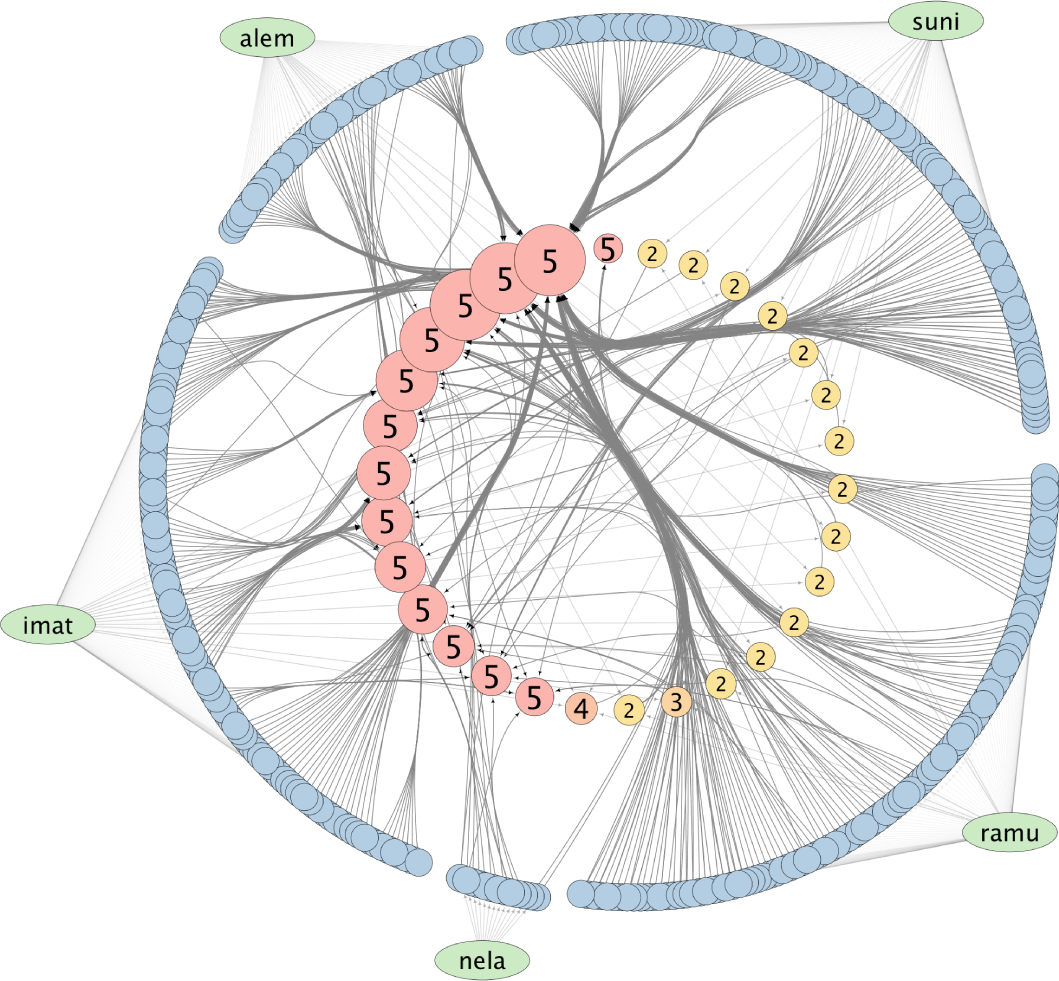
Core Publications in Networks. The arcs of blue nodes identify first generation publications (citing_sid) for each therapeutic. Nodes in the inner ring are sized by a gradient proportion to total degree count with an upper limit of 30 and are colored by a gradient proportional to the number of drug connections (2 to 5). 14 publications are common to all five networks (Table 3) and are colored red. The remaining nodes in the inner ring connect to between 2 and 4 drugs each and are labeled accordingly. Abbreviations: alem (Alemtuzumab), imat (Imatinib), nela (Nelarabine), ramu (Ramucirumab), suni(Sunitinib).

### Grant Support

With its annual budget of approximately US$32 billion, NIH is a major funder of biomedical research through its granting program. Understanding the nature and extent of NIH grant support for the research represented in our five networks, provides insight into the funding programs that enabled this research. We took advantage of publicly available data [12] to identify grant support for the publications in our five networks by mapping them to pmids. A total 19,104 unique grant numbers was harvested of which 112 were found in all five networks. At the intersection of five networks, the reason the number of grants is larger than the number of publications is because publications and grants exist in a ‘many to many’ relationship in that each publication can acknowledge support from multiple grants and each grant can support multiple publications. These awards were grouped by major type and visualized (Fig 3.). Of note, support from Research Program Projects and Center grants is proportionately larger in the intersection group when compared to the total population where the proportion of research projects is larger. A significant loss of information occurs when mapping from cited_sid to cited_pmid (Table 2). Thus we believe that these numbers may be an underestimate of actual grant support from NIH. Also missing from this analysis are details of research support from other funding agencies and industry, which are questions that we intend to pursue. Even so, these data testify to a recurring theme of collaboration and breadth of community engagement that is also seen at the publication level. We speculate that the broader and collaborative nature of such awards may be more likely to result in a methods-rich population of publications than the more focused research project award but elucidation will require further and more rigorous study.

**Fig 3.**
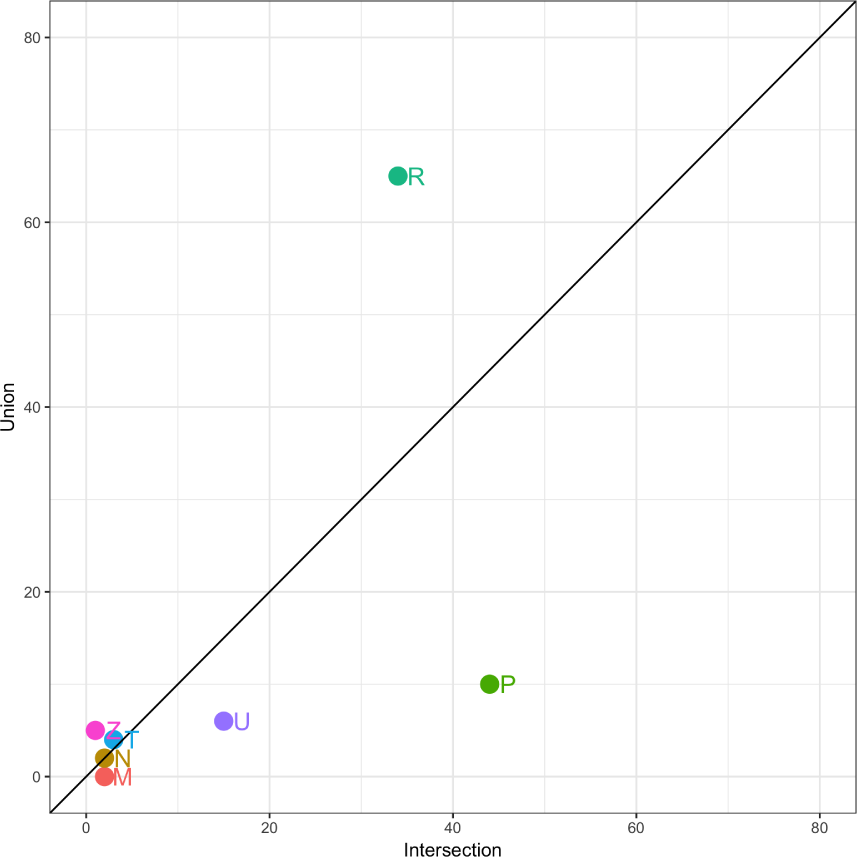
NIH Research Support. Grant and contract support for publications from NIH in the five networks was identified using ExPORTER data (Materials and Methods). 19,104 unique project numbers were identified as sources of support for publications in all five networks. Of these, 112 projects were common to all five networks. Projects were grouped by mechanism (i) P-Research Program Projects and Centers (ii) R-Research Projects (iii) M-General Clinical Research Centers Programs (iv) N-Research and Development-Related Contracts (v) U-Cooperative Agreements (vi) T-Training Programs (vii) Z-Intramural Research. For each mechanism, the number of projects in the intersection of all five networks was plotted against the number in the union of all five networks (both expressed as percentages of their respective totals). A higher proportion of Research Program Projects and Centers awards is found in the intersection group.

### Peer Review

Research support from NIH is typically made through a two stage peer-review process. The Center for Scientific Review at NIH manages first-stage scientific review of between 50,000-60,000 grant applications each year [19], a process involving more than 15,000 expert reviewers. In addition, individual Institutes and Centers at NIH manage smaller scale peer review operations. Considering a crude estimate of a 20% success rate in funding, peer review can be viewed as a collaborative scientific activity and that serves as a selection layer for the upper fifth of applications, thus strongly influencing granting outcomes. To describe this layer at a high-level, we matched the awards in the the five networks to the peer review panels (study sections) that evaluated them for scientific merit and calculated the intersection and union of these peer review panels. Eighty eight unique panel identifiers formed the intersection. Of these, 11 are distinguishable as Special Emphasis Panels that could be either one-time or recurring panels with temporary members, the remaining 77 are chartered panels with relatively stable membership. Some of these panels are no longer active and public records are not easily available to determine their scientific focus. For the 74 panels that could be classified (Supplemental), beyond an obvious focus on cancer, it is evident that the panels represent a rich mix of disciplines such as chemistry, biophysics, genetics, cell biology, and molecular biology; as well as AIDS, pathology, radiology, endocrinology, neurology, mental health, and child health. Four hundred and seven unique panel identifiers formed the union of all five networks. Of these 28 were Special Emphasis Panels, the remaining 379 panels were chartered as in the case of the intersection. These data provide evidence of broad input from invited experts in a collaborative activity that selects promising scientific projects. Assuming an average of 25 reviewers per panel (the number is likely to range from 5-40) and excluding that some of these panels are likely to have met multiple times during the lifespan of the awards in question and that some of these applications for funding may have been reviewed multiple times, a minimum of 10,000 experts comprised this additional layer of scientific influence. We believe that the actual number is likely to be at least double. A more accurate estimate would be possible if historical records of participation in peer review were made publicly available by NIH. We do not have records of awards or peer review from sources other than NIH and this also is a focus of future investigation.

### Network Metrics

To quantify network data and to identify influential researchers in and across networks, we calculated PIR and RBR scores for all researchers as well as nPIR, PPR, nRBR, and gRBR scores (Materials and Methods, Supplemental Information, Table 2). The nPIR metric describes the sum of aggregated citation scores for all articles that a researcher has in all five networks. The PIRpartitionRatio (PPR) results from normalizing nPIR to the sum of individual PIR scores to gain insight into holo-network influence. A limitation of the nPIR and nRBR measurements is that they are valid only for the network(s) being studied. Scaling from five to the more than 1400 drugs approved by the FDA (and their many variants) would address this limitation [20] although other data related issues may well emerge.

While theoretically appealing (Materials and Methods), the gRBR is the most sensitive to data quality since it relies on an accurate estimate of total productivity of a researcher, which is sensitive to data quality in bibliometric databases. For example, we found several instances in the top 10% of PIR scores where the gRBR was implausible likely on account of polysemy, synonymy, or incompleteness. This metric is therefore likely to be useful when the author disambiguation problem and article capture is resolved to the point where data quality is significantly improved and is not recommended except when strong confidence exists in the total productivity counts. These metrics may be best used in conjunction with positional measures such as quantiles to define populations of researchers within related networks, e.g, the top 25 researchers based on nPIR scores of all researchers in our dataset (S3 Table 2). These ‘bright stars in neighboring constellations’, represent elite performers in network(s) of clinical and basic science expertise again reinforcing the concept of collaborative translational achievements built upon a body of basic science. Beyond simple aggregation, weighting, and normalization that we have used, a variety of citation metrics such as SNIP [21], with different normalization strategies at the field, journal, and article level are available for impact analysis and these could be applied to such networks depending on the features of these networks and the aim of the study [22]. The use of these citation measures will assume greater importance when scale up from small numbers of networks to a greater proportion of the global network.

In summary, we have demonstrated a digital methodology based on data mining and network analysis, not restricted to drug discovery and cures alone, that offers burden-reduction in explorations of science history. Beyond assembling a set of facts about a major scientific advance, it contributes to the understanding of collaboration across domains and can be used to enrich portfolio analysis, planning and optimization, as well as communications of the societal value of research. The approach can be easily adapted to study the collaborative history within and across research portfolios of groups of researchers and targeted programs such as the Clinical and Translational Science Award (CTSA) Program. While finer critical evaluation of the content of datasets generated through this approach is best left to experts, the methodology is broadly accessible and can also be viewed as another tool for citizen science. Overall, no single metric will provide useful answers, instead expert interpretation of multiple metrics best matched to curated datasets will be valuable.

## Acknowledgments

This study would not have been possible without data that is made publicly available by NIH and the FDA. We thank Tandy Warnow for critical comments and advice on formalizing definitions. We thank Sydney Gomes and Caitlin Werle for assistance with graphics. We thank Sandeep Somaiya from NETE as well as Daniel Calto and Holly J. Falk-Krzesinski from Elsevier for their support of this project.

## Supporting Information

**S1 File. Network Calculations** The basis of calculations for network metrics.

**S2 Table 1 Intersecting Publications Across Networks.**

**S3 Table 2 Elite Performers in Networks**

